# Unveiling the Immune Dynamics of Neisseria Persistent Oral Colonization: A Roadmap for Innovative Vaccine Strategies

**DOI:** 10.1101/2023.12.18.572139

**Authors:** Mario Alles, Manuja Gunasena, Tauqir Zia, Adonis D’Mello, Saroj Bhattarai, Will Mulhern, Luke Terry, Trenton Scherger, Saranga Wijeratne, Sachleen Singh, Asela J. Wijeratne, Dhanuja Kasturiratna, Hervé Tettelin, Nathan Weyand, Namal P.M. Liyanage

**Affiliations:** Department of Microbial Infection and Immunity, College of Medicine, Ohio State University, Columbus, OH, USA; Department of Veterinary Biosciences, College of Veterinary Medicine, Ohio State University, Columbus, OH, USA; Infectious Diseases Institute, The Ohio State University, Columbus, OH, USA; Department of Biological Science, Ohio University, Athens, OH USA; Department of Microbiology and Immunology, Institute for Genome Sciences, University of Maryland School of Medicine, Baltimore, MD, USA; Institute for Genomic Medicine, Nationwide Children’s Hospital, Columbus, OH, USA; Arkansas Biosciences Institute, Arkansas State University, AR, USA; Department of Mathematics and Statistics, Northern Kentucky University, KY, USA; The Infectious and Tropical Disease Institute, Ohio University, Athens, OH, USA; Molecular and Cellular Biology Program, Ohio University, Athens, OH, USA

**Author notes:** Corresponding author: Namal P.M. Liyanage Assistant Professor, 788 Biomedical Research Tower, 460 W 12^th^ Ave, Columbus OH 43210. Equally contributed (Mario Alles, Manuja Gunasena).

## Abstract

Commensal bacteria are crucial in maintaining host physiological homeostasis, immune system development, and protection against pathogens. Despite their significance, the factors influencing persistent bacterial colonization and their impact on the host still need to be fully understood. Animal models have served as valuable tools to investigate these interactions, but most have limitations. The bacterial genus *Neisseria*, which includes both commensal and pathogenic species, has been studied from a pathogenicity to humans’ perspective, but lacks models that study immune responses in the context of long-term persistence. *Neisseria musculi*, a recently described natural commensal of mice, offers a unique opportunity to study long-term host-commensal interactions. In this study, for the first time we have used this model to study the transcriptional, phenotypic, and functional dynamics of immune cell signatures in the mucosal and systemic tissue of mice in response to *Neisseria musculi* colonization. We found key genes and pathways vital for immune homeostasis in palate tissue, validated by flow cytometry of immune cells from lung, blood and spleen. This study offers a novel avenue for advancing our understanding of host-bacteria dynamics and may provide a platform for developing efficacious interventions against mucosal persistence by pathogenic *Neisseria*.

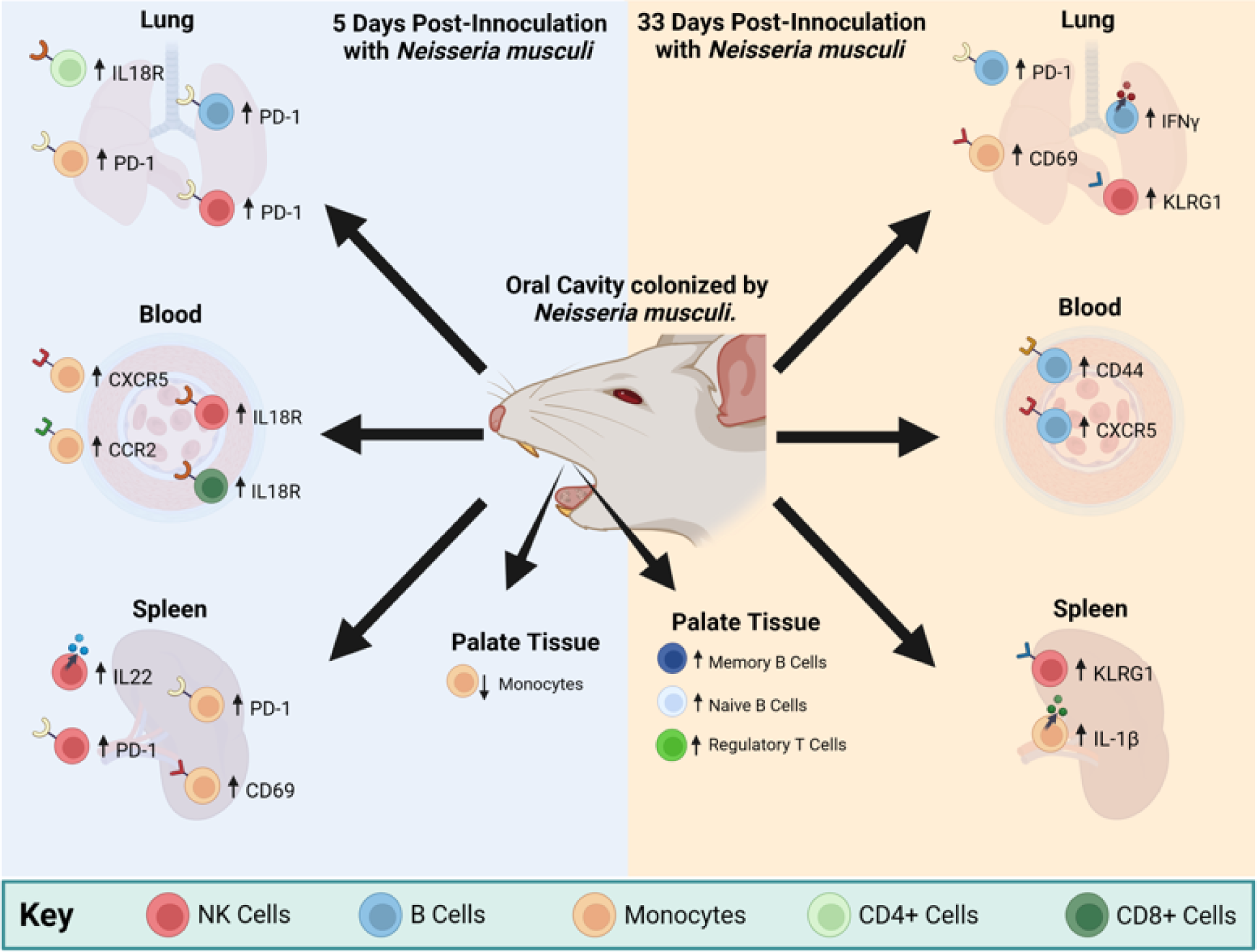

## Introduction

Commensal bacteria are an important component of a host’s physiological environment and play a critical role in its functional homeostasis, immune system development and protection against pathogens^1,2^. Despite their importance, the factors influencing persistent bacterial colonization and its impact on the host’s immune system, remain understudied. To delve deeper into commensal-host interactions and their consequences, animal models have traditionally served as valuable tools.

*Neisseria* has been investigated for its commensal nature within mammalian hosts. It encompasses many species that are genetically related^3^. These can colonize a broad range of hosts ranging from rodents to non-human primates to humans^4^. The two primary human pathogens, *N. gonorrhoeae* and *N. meningitidis*, are associated with high levels of asymptomatic colonization and can straddle the border between commensalism and pathogenicity^5^. Given the absence of an effective vaccine for gonococci, various infection models have been created to enhance our comprehension of pathogen-host dynamics and uncover insights into the immune response and potential vaccine-induced protection mechanisms. Indeed, several mouse models have been proposed to study *N. gonorrhoeae* genital tract infections but due to its tropism for humans these models require several external manipulations to maintain transitory infections and therefore may not mimic natural conditions in which host-pathogen interactions can be fully assessed^6^. Pharyngeal infection models are limited; yet this site facilitates persistent colonization by both pathogenic *Neisseria* species for several months^7,8^. We have recently developed a mouse model to explore *Neisseria* colonization, using the newly characterized mouse commensal *Neisseria musculi*^9^. Retaining crucial host interaction genes in *N. musculi*, shared with *N. gonorrhoeae*, makes it an excellent surrogate species for modeling gonococcal pharyngeal colonization^9-11^.

In this study, we found multiple changes in the transcriptional profile of palate tissue colonized with *N. musculi* and the enrichment of the gene pathways associated with diverse biological processes. For the first time, we have unraveled the dynamic alterations in both innate and adaptive immune signatures within systemic tissue of colonized mice, fostering the establishment and maintenance of immune homeostasis. Our findings may advance the knowledge toward developing potential vaccine candidates efficacious against mucosal carriage of pathogenic *Neisseria* species.

## Results

### Oropharyngeal colonization by *Neisseria musculi* was established following oral inoculation of the bacterium

A total of 16 A/J mice were divided into two groups of eight mice each consisting of an inoculated group and control group. A slow oral inoculation of 50μL of *N. musculi* bacterial suspension was administered to each mouse in the inoculated group while control mice were mock inoculated with 50μL of PBS into the oral cavity (Figure 1A). Here we found that following inoculation the bioburden of *N. musculi* in colonized tissues of the tongue and hard palate were higher at day 33 (late) compared with day 5 (early) (Figure 1B). However, our serial monitoring of colony counts obtained through weekly sampling of oral swabs (Figure 1C) and fecal pellets (Figure 1D) suggest that bacterial colonies continue to increase and peak around the 3^rd^ week after inoculation with no further increases, suggesting a degree of homeostasis and tolerance between host tissue and commensal bacteria had been reached.

**Figure 1.**
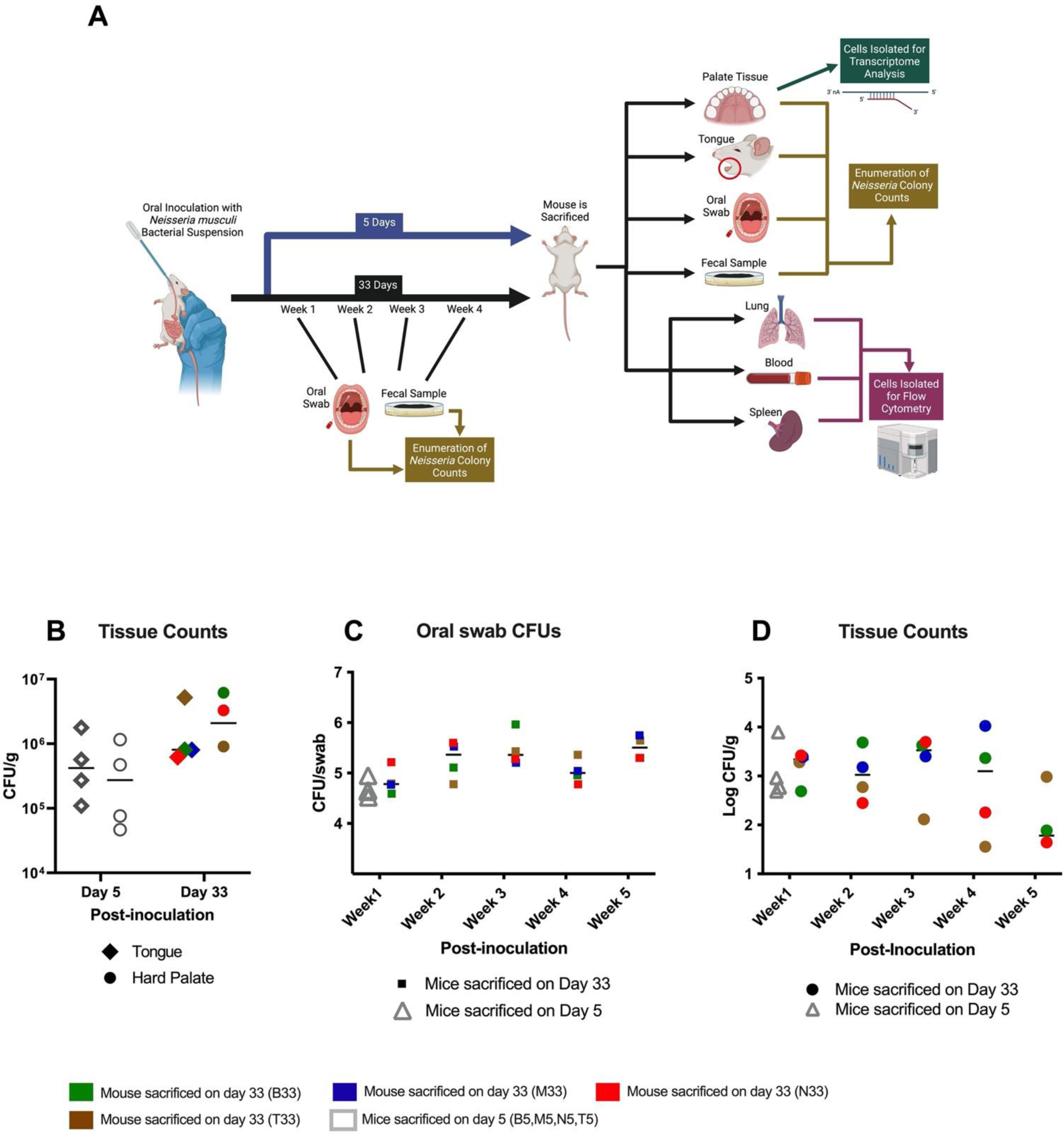
Experimental design and establishment of colonization following oral inoculation with *N. musculi*. **(A)** Mice from the experimental group (n=8) were orally inoculated with *N. musculi* followed by a subset (n=4) being euthanized after 5 days. Inoculated mice left for 33 days (n=4) before euthanasia were sampled weekly for enumeration of oral and fecal colony counts. Blood, lung and spleen were harvested at sacrifice for flowcytometric analysis of immune signatures between inoculated and control mice. Enumeration of oral colonization was done using colony counts of tongue and palate. Bulk RNA-seq was performed on palate tissue to observe differences in tissue transcriptome following colonization. **(B)** Dot plot depicting differential abundance of *N. musculi* colonies in colonized tongue and hard palate tissue at days 5 and 33 after inoculation. Dot plots represent weekly changes in *N. musculi* colony counts obtained from culture of **(C)** oral swabs and **(D)** fecal pellets of inoculated mice. CFU/g, colony forming units per gram of tissue. Animal M33 did not have palate colony counts and fecal counts for week 5.

### Transcriptome analysis reveals dynamic changes in gene expression and immune cell composition during *Neisseria musculi* colonization in palate tissue

We performed bulk RNA sequencing (RNA-seq) on palate tissue of mice inoculated with *Neisseria musculi* and then sacrificed on day 05 and day 33 post-inoculation, as well as their control mice. Based on the expression of selected biomarkers from our transcriptome analysis, we observed that host gene expression patterns distinguish mice colonized with *N. musculi* from their controls (Supplementary Figure 1). While no major differences in gene expression were noted by day 5 after inoculation, at day 33 a total of 26 differentially expressed genes (DEGs), including 20 upregulated DEGs and 6 downregulated DEGs (Supplementary Table 1) were identified in the tissues of the palate of mice colonized with *N. musculi* compared with those of control mice (Figure 2A). Here we observed several DEGs linked to inflammation (*Ltb, Csf3, Tnf, Slamf7*), and mucosal immunity (*IL22ra, Igha*), likely attributed to colonization with *N. musculi*.

**Figure 2.**
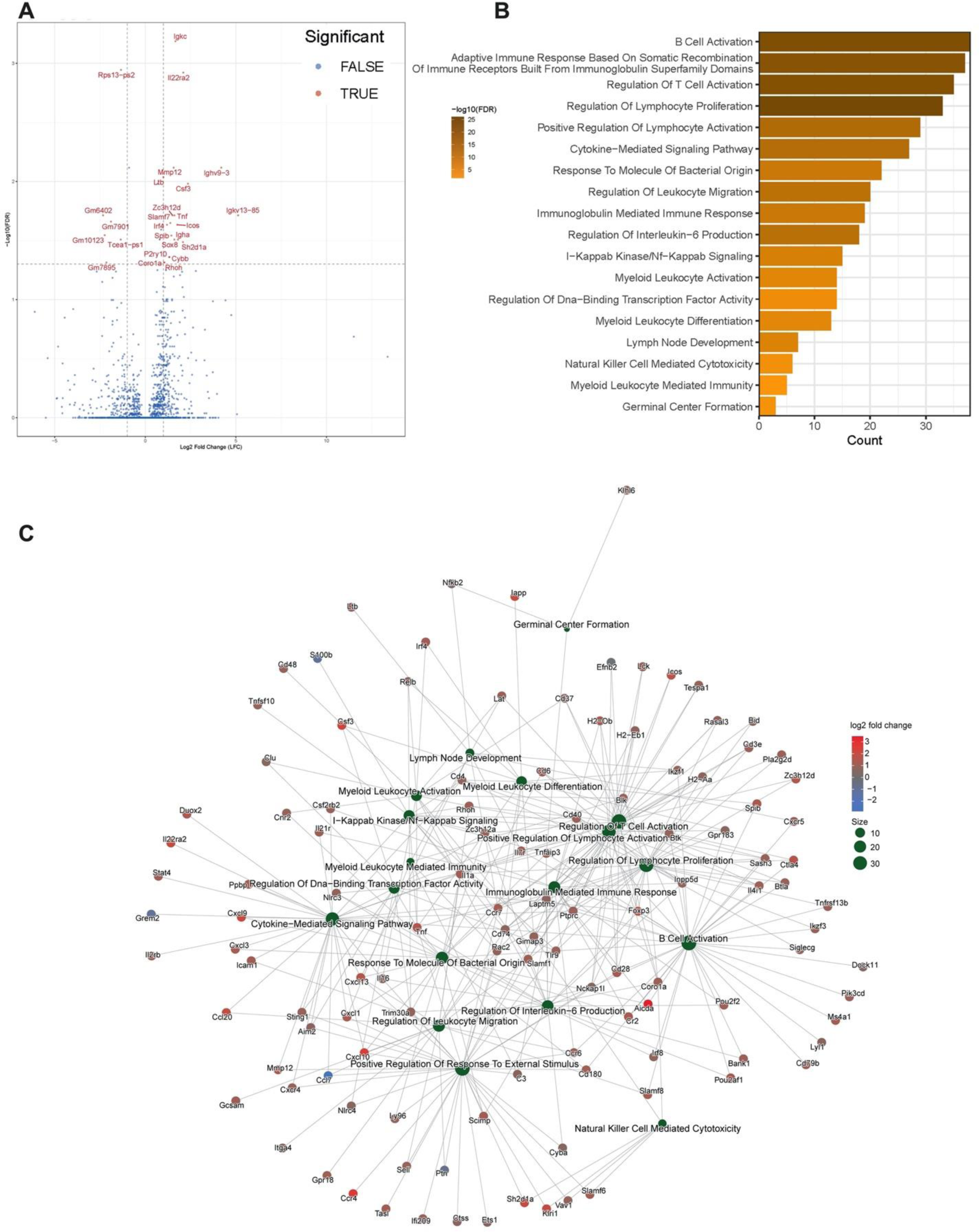
Differentially expressed genes (DEGs) and enriched transcriptional pathways of cellular biological processes in palates of mice 33 days after being orally inoculated with *N. musculi* vs mock-inoculated controls. (**A**) Volcano plot of DEGs between inoculated mice (n=4) and controls (n=4) (screening thresholds: adjusted p value < 0.05). **(B)** Gene Ontology (GO) analysis was performed and revealed significantly enriched pathways in palate tissue from inoculated mice vs. controls related to biological processes (screening threshold: adjusted p value < 0.05). **(C)** Cnet plot of GO biological process enrichment of DEGs of palate tissue of inoculated vs control mice shows distinct connections between biological processes resulting from colonization with *N. musculi*.

To analyze the biological importance of these DEGs, Gene Ontology (GO) enrichment analysis was performed (Supplementary Table 2). Based on the enriched GO terms, we observed significantly enriched pathways involving the biological activity of several immune cells including T cells, B cells, cells of the myeloid lineage and natural killer (NK) cells (Figure 2B). Of particular interest were enriched pathways involving *myeloid leukocyte activation and differentiation*, *NK cell-mediated cytotoxicity*, *T and B cell activation*, *lymphocyte proliferation*, *immunoglobulin mediated immune response*, *IL-6 production*, *germinal center formation* and *response to bacteria*. We also created network visualizations of selected enriched GO pathways using Cnet plots (Figure 2C and Supplementary Figure 2). Connections between pathways in the network imply shared genes, suggesting possible crosstalk or cooperation in enriched biological processes particularly related to leukocyte immune activation due to colonization with *N. musculi*.

We also performed deconvolution of our bulk transcriptome data of palate tissue of mice inoculated with *N. musculi* and control mice using TIMER2.0^12^. Using six algorithms we generated differential estimations of abundance of immune cell subsets in each sample of the two groups (Figure 3A-B and Supplementary Figure 3). Here, we found significant differences in the expression of immune cells at both time points (day 5 and day 33) following bacterial colonization of palate tissue (Figure 3C). At day 5 after inoculation, our data indicated significantly reduced expression of monocytes, the main producers of IL-6 which mediates partial resistance to bacterial colonization, in the palates of mice colonized with *N. musculi*. Conversely by day 33 after inoculation, data from deconvolution of the palate transcriptome indicated an overall enhancement of the adaptive immune response reflected by an altered tissue transcriptome suggesting increased expression of memory B cells, naïve B cells and regulatory T cells in the palates of mice colonized with *N. musculi* compared with palate tissue of controls. This suggests that with time, lack of resistance to *N. musculi* colonization establishment gives way to a more protective and regulatory immune response in colonized tissue.

**Figure 3.**
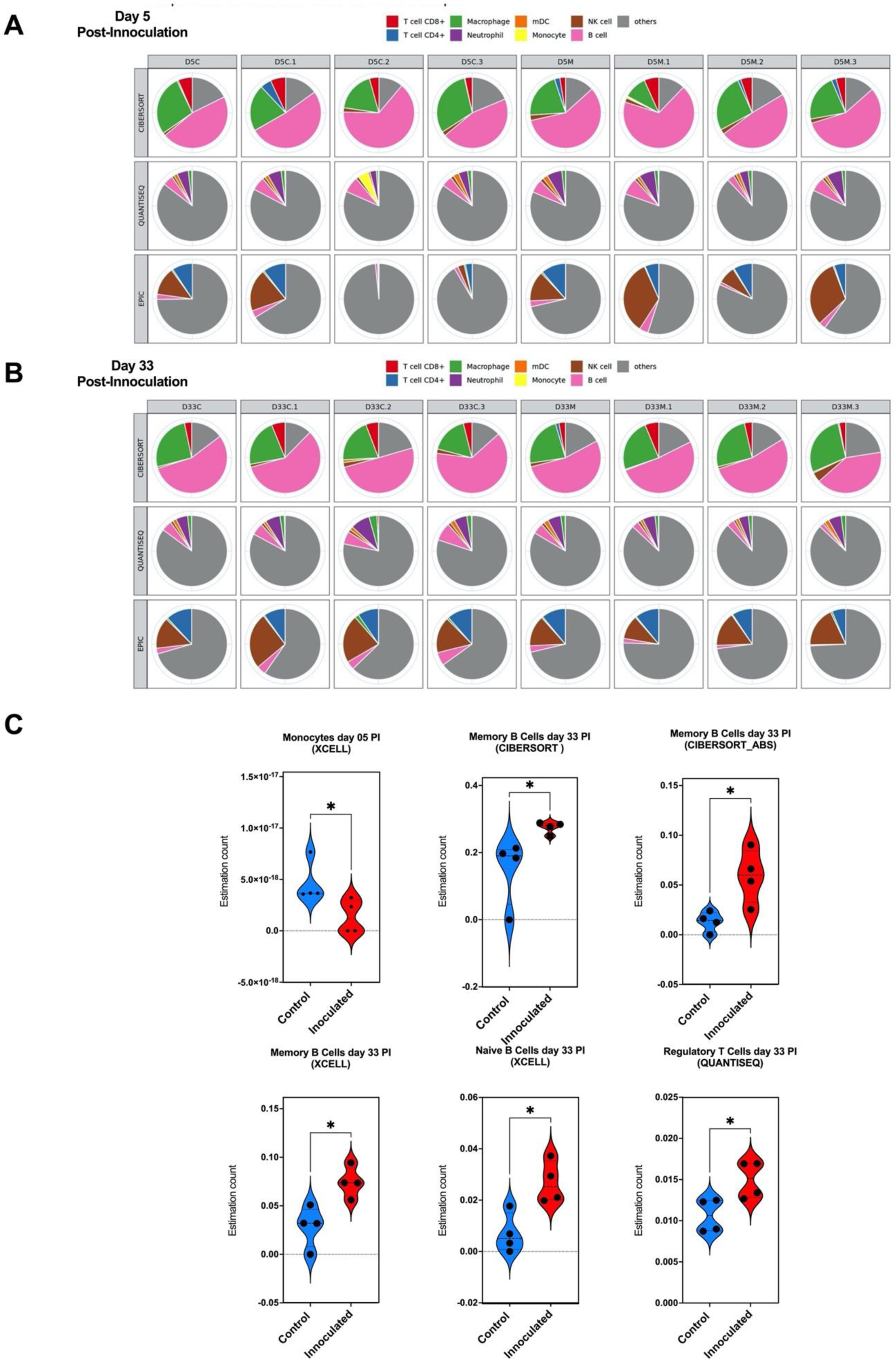
Deconvolution of transcriptome data from palate tissue of mice 5 and 33 days after being inoculated with *N. musculi* vs mock-inoculated control mice. The TIMER2.0 database was used to generate estimations of abundance of immune cell subsets in palate tissue of each mouse belonging to both time points using six algorithms. Multi-panel pie plots for three algorithms which generate comparable values show the proportions of major immune cell types in each sample at **(A)** day 5 and **(B)** day 33 after inoculation (including those of controls). D5C/D33C, mice colonized with *N. musculi* sacrificed on day 5 / day 33. D5M/D33M, mock-inoculated mice sacrificed at day 5 / day 33. **(C)** Violin plots based on algorithm-generated abundance scores depict differential expression of selected immune cell subsets at day 5 and day 33 after inoculation compared with control mice in corresponding time points. The p value was calculated using Wilcoxon rank-sum test *p<0.05. PI, Post-Inoculation.

### Dynamic and temporal changes are observed in the immune response of mucosal tissue that is anatomically adjacent to tissue colonized with *N. musculi*

Of the tissues analyzed by flow cytometry, we considered tissue of the respiratory system (i.e., lung) as mucosal tissue anatomically most adjacent to the inoculated (and therefore colonized) oral cavity^13^. We initially performed unbiased clustering of lung immune cells at both day 5 (Figure 4A) and day 33 (Figure 4B) after inoculation by utilizing FlowSOM-based automatic clustering algorithms, and UMAP for dimensional reduction visualization. This analysis was conducted on CD45^+^ cells to discern changes in immune cell clusters between mice inoculated with *N. musculi* and controls. While no major changes in clustering were observed between these two groups on day 5 after inoculation (Figure 4A), we observed changes in monocyte- and T cell-clusters at day 33 (Figure 4B) after inoculation (late colonization) suggesting differences in phenotypic expression of these immune cells in the lung mucosal tissue influenced by *Neisseria* colonization. Principal component analysis (PCA) performed using flow cytometric data revealed distinct clustering between inoculated mice and control mice at both day 5 (Figure 4C) and day 33 (Figure 4D) after bacterial inoculation, suggesting that colonization with *N. musculi* drastically alters the immunophenotype and functionality of mucosal immune cells in adjacent tissue. To understand changes induced by bacterial colonization in specific immune cell subsets, we performed manual gating (Supplementary Figure 4) of our flow cytometry data (Figure 4E). Many of the immune signatures significantly altered by bacterial colonization were dynamic and temporal, in that these alterations were limited to occur either early on (day 5) or later (day 33) during colonization. Interestingly we observed upregulation of PD-1 on B cells, which was consistently elevated in the lung tissue of inoculated mice throughout colonization. Given that these are unswitched, IgM-producing memory B cells, colonization therefore results in a persistently enhanced mucosal humoral immune response capable of generating antibodies on exposure to related antigens^14^. We also observed increased PD-1 expression in monocytes and NK cells of inoculated mice compared with controls, but only early on in colonization and reaching baseline later on day 33, suggesting tolerance and immune homeostasis developing following early persistent antigenic stimulation due to bacterial colonization^15^. Nevertheless, our results indicated that monocyte activity may still be enhanced with a more mature phenotype seen among NK cells even late into colonization as evidenced by increased CD69 expression in monocytes^16^ and increased KLRG1 expression in NK cells^17^, respectively. The enhanced activity of monocytes in lung tissue may be at least partially explained by significantly higher proportions of IL-18 receptor (IL18R) expressing CD4+ T cells (found to express more IFNγ)^18^ and IFNγ producing innate B cells in this tissue of colonized mice^19^ which also suggests a more pro-inflammatory environment occurring in lung mucosal tissue as a result of *N. musculi* colonization.

**Figure 4.**
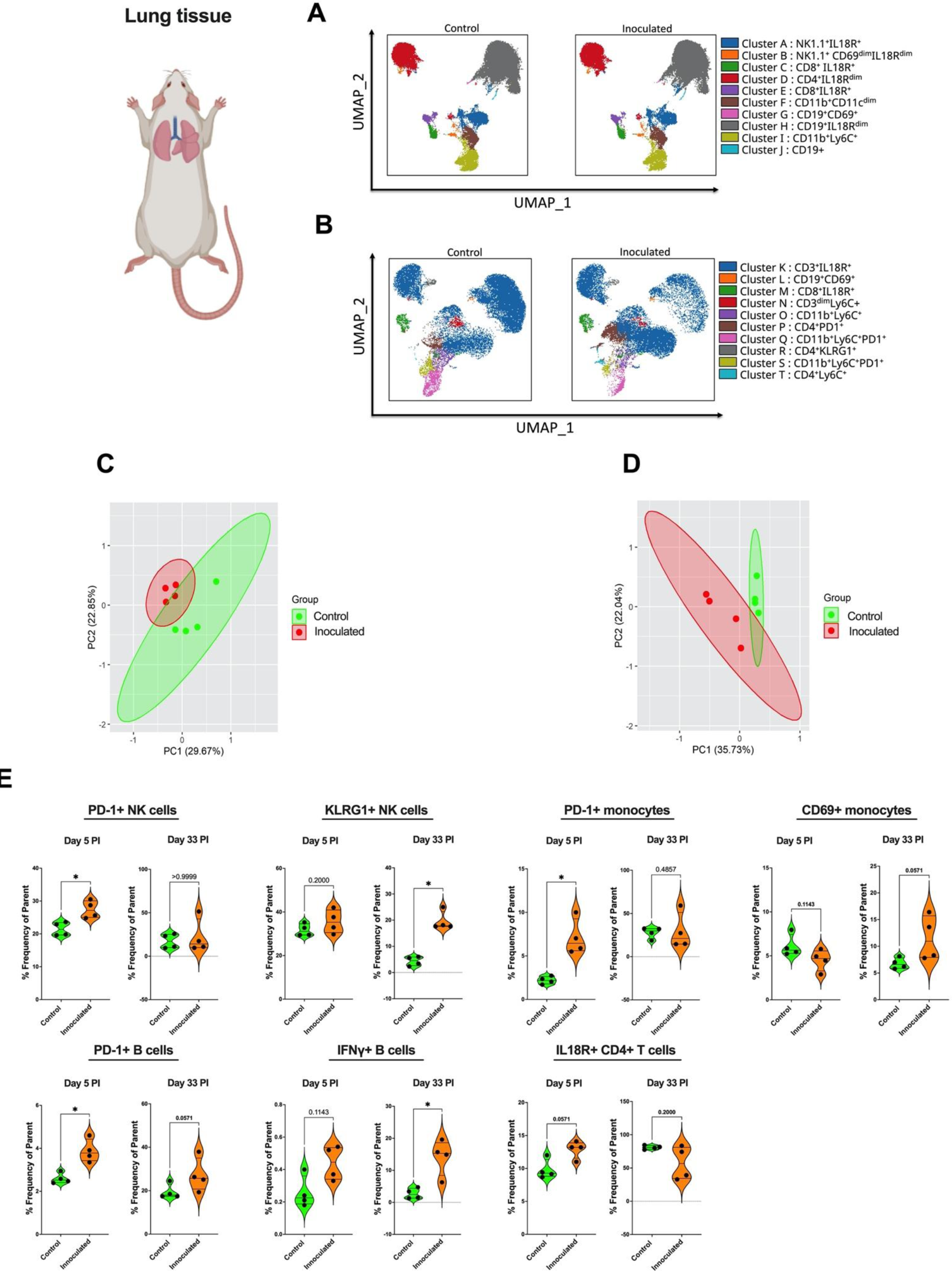
Alterations in the immune landscape of adjacent mucosal tissue in mice following oral colonization with *N. musculi*. FlowSOM-based clustering of CD45^+^ cells in mucosal lung tissue of mice **(A)** 5 days and **(B)** 33 days after inoculation with *N. musculi*, depicted on Uniform Manifold Approximation and Projections (UMAPs) reveal changes in phenotypic clusters compared with that of controls. Principal component analysis (PCA) was performed using selected immune markers and reveal distinct clustering of mice inoculated with *N. musculi* at **(C)** day 5 and **(D)** day 33, compared with controls. **(E)** Violin plots depicting differential expression of immune signatures between inoculated and control mice at day 5 and day 33 after inoculation. The p value was calculated using Wilcoxon rank-sum test *p<0.05. PI, Post-Inoculation.

### Oral inoculation with *N. musculi* alters the systemic immune response in mice towards a pro-inflammatory phenotype early on after exposure to the bacterium

Alterations in immune signatures in the blood and spleen were representative of the systemic immune response in mice following colonization with *N. musculi*. FlowSOM-based unbiased clustering visualized on UMAP revealed changes in expression of CD45+ cell clusters in both the blood and spleen in mice 5 days after bacterial inoculation versus controls. In terms of immune responses in the blood, we specifically observed differential expression of monocytes clusters and T cell clusters between the two groups (Figure 5A) whereas in the spleen changes were most apparent in monocyte clusters (Figure 5B). To further delineate more specific differences in immune cell subsets brought on by *N. musculi* oral inoculation, we performed manual gating of our flow cytometry data from immune cells of blood and spleen. Based on PCA, overall changes in immune cell markers of the blood revealed distinct clustering of mice inoculated with *N. musculi* compared to controls (Figure 5C). Detailed analysis of peripheral blood immune signatures revealed that colonization with *N. musculi* early on (day 5) increased the migratory capacity of blood monocytes to inflamed mucosal tissue and secondary lymphoid organs by increasing their expression of surface expression of CCR2^20^ and CXCR5^21^, respectively (Figure 5D). Furthermore, we found that oral inoculation resulted in peripheral cytotoxic immune cells including NK cells and CD8+ T cells displaying a more pro-inflammatory phenotype, as evidenced by their increased expression of IL-18R compared with controls, suggesting more IFNγ production by these cells^22,23^. In the spleen, we observed significant differences in immune phenotypes between mice inoculated with *N. musculi* compared to control mice (Figure 5E). Here, we found increased PD-1 expression among NK cells and monocytes following inoculation suggesting their response to persistent antigenic stimulation^24^, coupled with enhanced activation of monocytes suggested by increased CD69 expression. Interestingly, inoculation with *N. musculi* also increased IL-22 production by NK cells of the spleen, suggesting their role in maintaining mucosal immunity during the early phase of colonization (Figure 5F)^25^.

**Figure 5.**
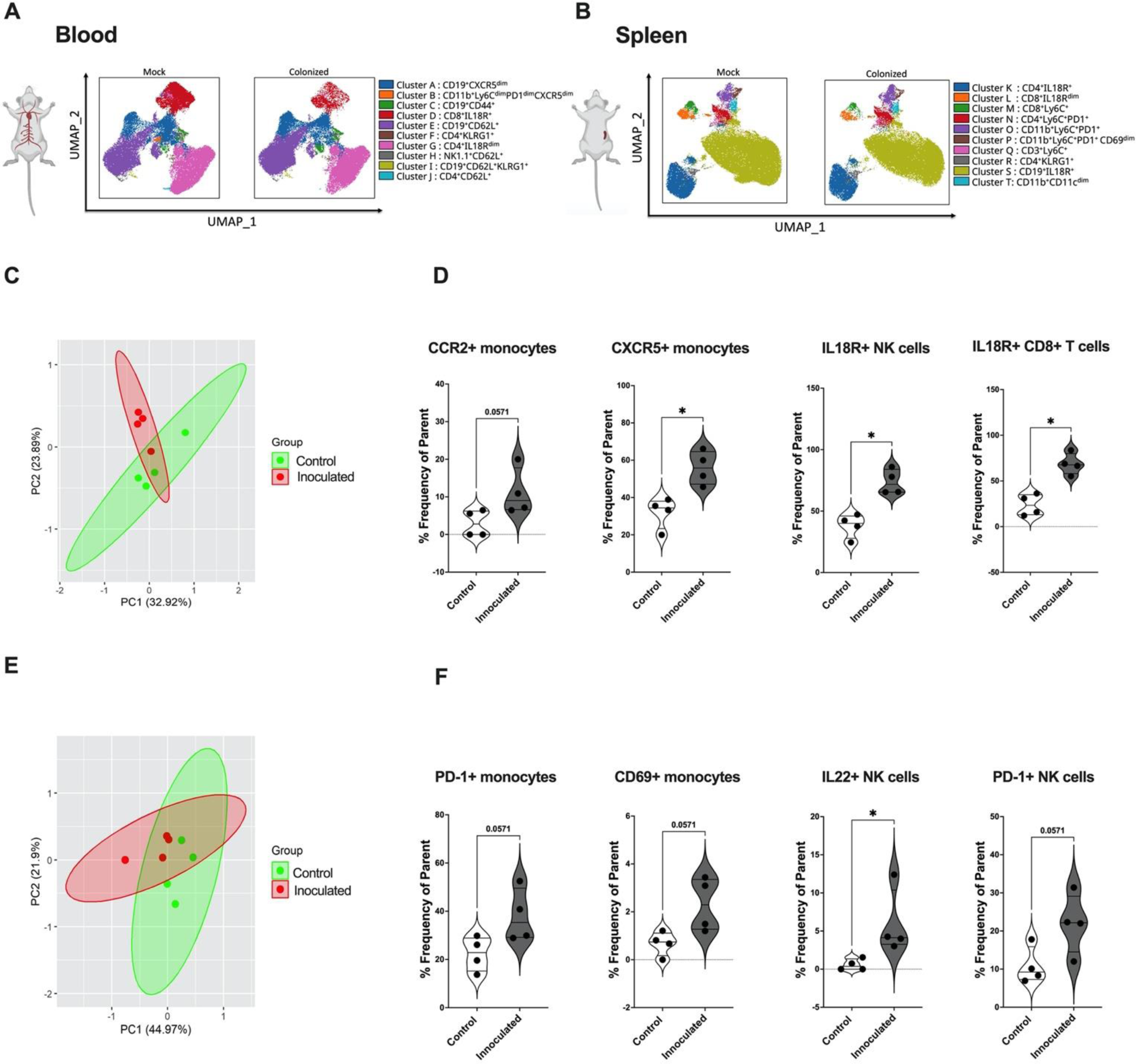
Alterations in the systemic immune landscape of mice early on following oral colonization with *N. musculi*. FlowSOM-based clustering of CD45^+^ cells in **(A)** blood and **(B)** spleen of mice 5 days after inoculation with *N. musculi*, depicted on Uniform Manifold Approximation and Projections (UMAPs) reveal changes in their phenotypic clusters compared with that of controls. Principal component analysis (PCA) was performed using selected immune markers in **(C)** blood and **(E)** spleen and reveal distinct clustering of mice inoculated with *N. musculi* at day 5 compared with controls. Violin plots depicting differential expression of immune signatures in **(D)** blood and **(F)** spleen between inoculated and control mice at day 5 after inoculation. The p value was calculated using Wilcoxon rank-sum test *p<0.05.

### Colonization with *N. musculi* maintains a pro-inflammatory immune environment within systemic tissue even towards the later stages following inoculation

Our immune data obtained from the blood at day 33 after inoculation with *N. musculi* indicated notable changes in the expression of B and T cell clusters (Figure 6A), whereas in the spleen, it was evident that monocytes and B cell clusters were the most prominently affected (Figure 6B). Based on PCA analysis, we also observed distinct immune responses in mice, at day 33 after *N. musculi*, colonization in the blood (Figure 6C) and spleen (Figure 6E). However, it is noteworthy that the prominent changes in the immunophenotype within both these tissues at day 33 post-inoculation (Figure 6D and F) were less frequent than those observed at day five post-inoculation. These findings suggest that over time, a certain degree of immune homeostasis and tolerance is established within these systemic tissues, supported by the patterns identified through PCA analysis. However, we still observed increased migratory capabilities of antigen primed immune cells, more specifically B cells in the blood towards secondary lymphoid tissue and inflamed tissue indicated by their increased expression of CXCR5 and CD44^26^. Additionally, immune data from spleen tissue at day 33 after inoculation with *N. musculi* revealed more activated pro-inflammatory immune cells indicated by increased expression of IL1β from monocytes compared with controls^27^. At the same stage of colonization splenic NK cells were found to be of a more mature phenotype indicated by their increased expression of KLRG1, further confirming the impact of *N. musculi* colonization on the innate immune compartment.

**Figure 6.**
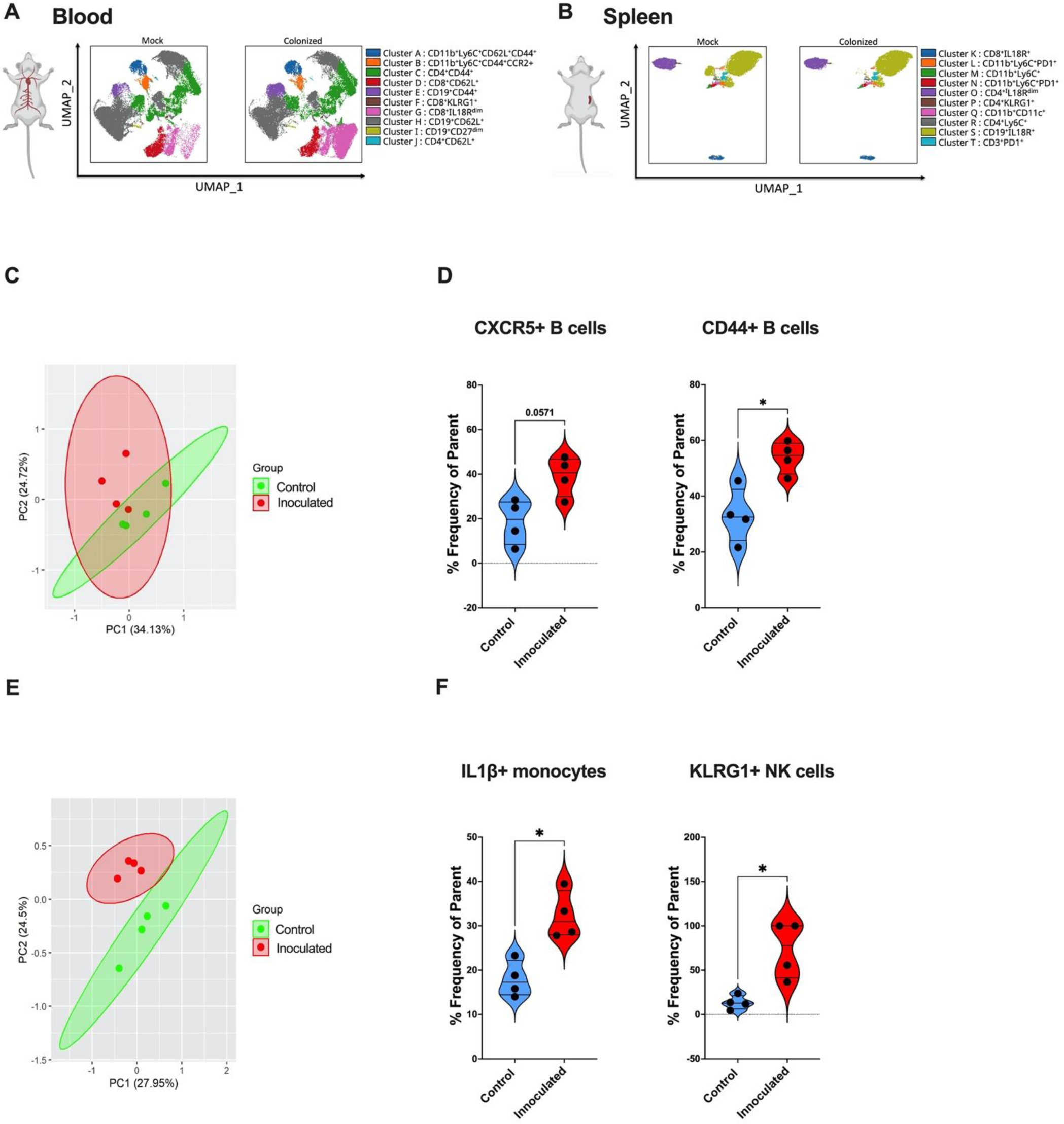
Alterations in the systemic immune landscape of mice in later stages following oral colonization with *N. musculi*. FlowSOM-based clustering of CD45^+^ cells in **(A)** blood and **(B)** spleen of mice 33 days after inoculation with *N. musculi*, depicted on Uniform Manifold Approximation and Projections (UMAPs) reveal changes in their phenotypic clusters compared with that of controls. Principal component analysis (PCA) was performed using selected immune markers in **(C)** blood and **(E)** spleen and reveal distinct clustering of mice inoculated with *N. musculi* at day 33 compared with controls. Violin plots depicting differential expression of immune signatures in **(D)** blood and **(F)** spleen between inoculated and control mice at day 33 after inoculation. The p value was calculated using Wilcoxon rank-sum test *p<0.05.

### *N. musculi* colonization is associated with PD-1 expressing immune cells

We performed Pearson correlation coefficient analyses between colony counts of *N. musculi* in palate tissue at both days 5 and 33, and corresponding immune cell subsets in tissues. This revealed that the magnitude of colonization of palate tissue correlated with several immune cell subsets expressing the immune checkpoint marker PD-1, including splenic NK cells (Figure 7A), splenic monocytes (Figure 7B) and B cells of the lung (Figure 7C). Given that PD-1 expression is associated with persistent antigenic stimulation and that the above immune subsets were found to be increasingly expressed at one or both time points following inoculation, this indicates that *N. musculi* directly influences the systemic and mucosal tissue immune landscape of the host.

**Figure 7.**
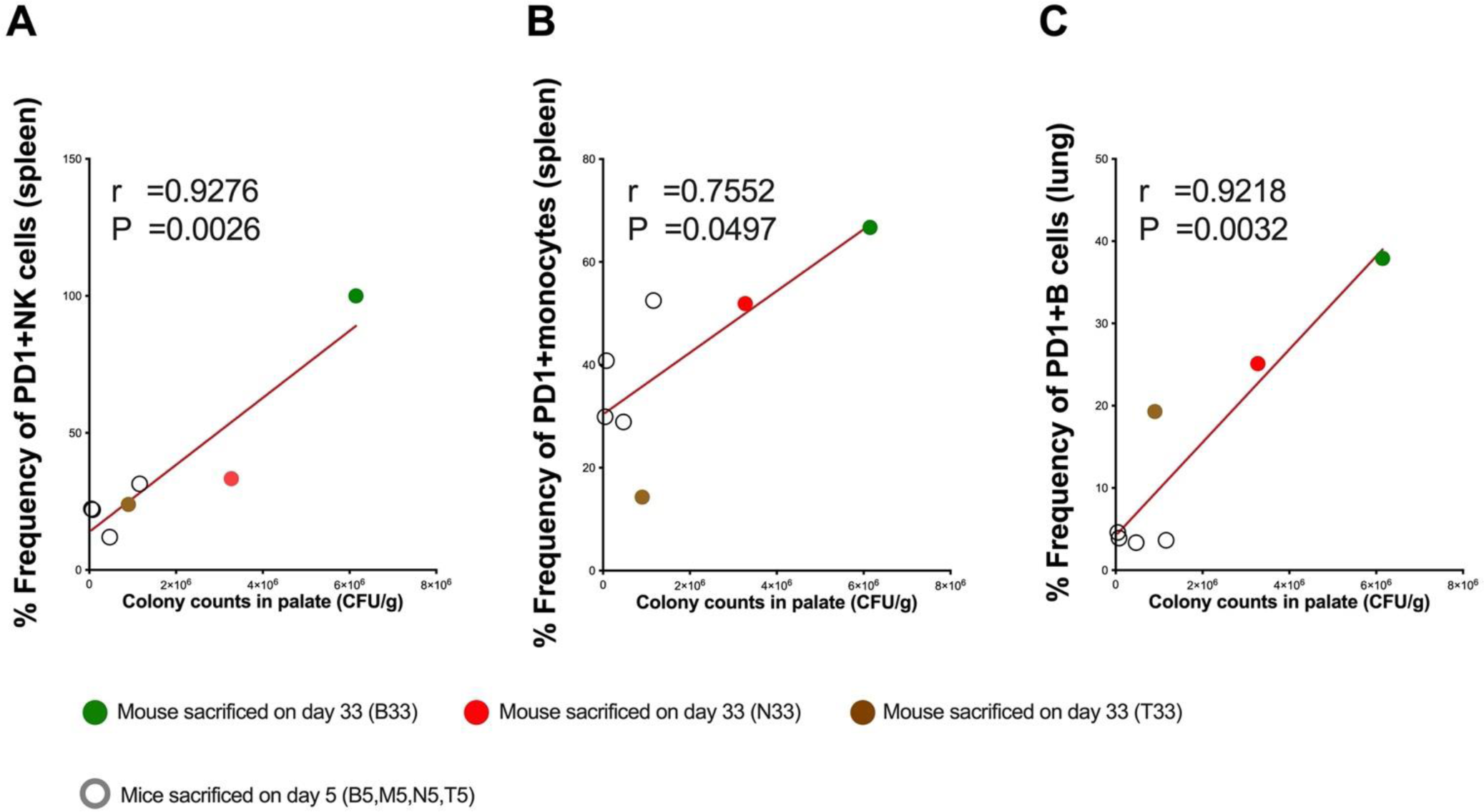
Correlational analysis between *N. musculi* colonization and immune signatures. Pearson correlation coefficient analysis revealing correlations between magnitude of colonization of *N. musculi* in palate and expression of PD-1 in **(A)** splenic NK cells, **(B)** splenic monocytes and **(C)** B cells in lung. r, Pearson correlation coefficient. P, p value.

## Discussion

The primary focus of *Neisseria* colonization studies has been on the human pathogens, *N. gonorrhoeae* and *N. meningitidis*, with a goal to identify targets for vaccine antigens that can induce protective immune responses against these pathogens^28,29^. Many homologs of pathogenic *Neisseria*-host interaction factors and candidate vaccine antigens have been identified in the mouse commensal *N. musculi*. Our study unveiled notable changes in the expression of both systemic and mucosal innate and adaptive immune signatures following oral inoculation with *N*. *musculi*. As demonstrated by Powell et al., cytokines like IL-6 are pivotal in conferring partial resistance against *N. musculi* colonization^30^. Notably, monocytes stand out as the principal immune cells responsible for IL-6 secretion^31^. Interestingly, in this study, we observed reduced expression of monocytes signatures in the locally colonized palate tissue revealed by our transcriptomic deconvolution data in early colonization. Furthermore, at this early-stage colonization, we also noted increased expression of monocytes with upregulated chemokine receptors (i.e., CCR2) and activation markers in both the blood and the spleen. Therefore, we suggest that the innate immune response promotes local tissue colonization and simultaneously walls of the colonized tissue and limits systemic spread.

Additionally, we observed increased systemic cytokine responses in early colonization, including upregulation of IL-22 production by splenic NK cells, a crucial cytokine involved in maintaining epithelial integrity and promoting bacterial colonization in lymphoid tissue, contributing to more robust systemic immune responses^32^. We also have observed similarities in our transcriptomic data where we found upregulation of *IL22ra* gene expression in palate tissue during late colonization, suggesting a systemic response geared to regulate colonization of *N. musculi* and promote a lymphoid tissue response to the bacteria. Our data also suggest an overall pro-inflammatory response taking place in the early phase of colonization indicated by increased expression of peripheral blood T cells expressing IL18R which in turn are known to produce high levels of IFNγ^18^. Similar to the discoveries by Baldridge et al.^33^, wherein they revealed that IFNγ initiates the activation of dormant hematopoietic stem cells within the backdrop of chronic infection, we propose a similar phenomenon occurring during early colonization by *N. musculi*. Here increased systemic IFNγ response could promote hematopoietic stem cell activation and enhance the immune response. Thus, we propose a local and systemic immune response occurring in the early phase of colonization geared towards promoting the establishment of *N. musculi* colonization in local inoculated tissue while the systemic immune response also builds up its defenses to ensure the host is protected from potential systemic effects of the bacteria.

The immune response we observed during the late colonization differs significantly from the early phase of colonization. During the later stages of colonization (i.e., day 33 after inoculation) our specific emphasis was directed toward examining the immune response within the mouse lung. Given its anatomical adjacency to the colonized palate, we identified the lung as an instrumental system profoundly influenced by *N. musculi* colonization. Hence, we deem the lung as a surrogate marker, providing insights into the immune response observed not only in mucosal tissues but also in anatomically neighboring sites affected by colonization. Given that pathogens like *N. gonorrhoeae* also colonize mucosal tissues of the upper respiratory tract, the data we gather by analyzing these tissue types will likely be invaluable during future vaccine development. The proposed link between colonized palate tissue and lung tissue was partially confirmed by deconvoluting transcriptome data from the palate, which showed similar signature changes to those observed in the late stages of lung tissue colonization. Here we found the immune response to be protective and geared up to defend against further bacterial invasion. In the monocytes of lung tissue, we found increased expression of activation markers such as CD69, signifying increased production of IL-6 to limit the spread of bacterial colonization (as described earlier). Enrichment of GO terms corresponding to myeloid cell activation and differentiation further confirmed our results. A higher population of unswitched, IgM-producing memory B cells, signified by their increased expression of PD-1^14^, and confirmed by enrichment of GO terms corresponding to the humoral response means that colonization likely stimulates local neutralizing antibody production against the offending antigen. In late colonization, an increase in IFNγ-producing B cells suggests the sustained presence of a pro-inflammatory environment within this tissue. We also observed increased expression of KLRG1 on the NK cells of lung tissue suggesting that even innate immune cells in this tissue adjacent to colonized regions display a more mature phenotype and also contribute to immune homeostasis^34^. This was likely primed by signals generated from adjacent colonized tissue, the transcriptome data of which showed enrichment of related pathways like leukocyte differentiation. We also observed upregulated immune markers associated with increased maturity (KLRG1+ splenic NK cells), pro-inflammatory activity (IL1β secreting monocytes in blood), and lymphoid tissue homing capability (CXCR5 expressing B cells in blood) in the systemic immune compartment even late into colonization. What is interesting to note is that even late into colonization, these changes in immune signatures persist in these tissues. The significant correlations we observed between the magnitude of colonization with *N. musculi* in palate tissue and the expression of the immune marker PD-1 in many systemic immune cells further confirms the influence of local bacterial colonization on the systemic immunophenotype. While the phenotypic and functional attributes of PD-1 in conventional T cells are well established^35^, less is known about the significance of PD-1 expression in other immune cell types. Indeed, from a T cell perspective, PD-1 is found to be expressed during activation and shows high and sustained expression during persistent encounters with antigen, much like during persistent bacterial colonization. The PD-1 pathway limits overactivation during initial priming and fine-tunes effector cell differentiation when exposed to antigen. Furthermore, in tissues such as the lung during acute infection, the PD-1 pathway protects tissues from immunopathology, regulates memory cell formation, and mediates a return to immune homeostasis^15^. Limited studies on PD-1 expression on monocytes confirm that elevated levels of microbial products and inflammatory cytokines in the blood cause upregulation of PD-1 on monocytes. Furthermore, it has been shown that PD-1 expression on monocytes correlated with IL-10 concentrations in the blood^36^. IL-10 has potent anti-inflammatory properties and plays a key role in limiting host immune response to microbes, thereby preventing damage to the host, and maintaining normal tissue (particularly mucosal) homeostasis. While we have already described the importance of maintaining immune homeostasis during bacterial colonization, further research is needed to delineate the more specific roles of the PD-1 pathways in the other immune cell subsets.

Here we have developed a murine model for investigating immunological determinants of *N. musculi* colonization and persistence. Its similarities with human pathogenic *Neisseria* species make *N. musculi* and ideal candidate species to investigate the transcriptional and immunological events surrounding long-term persistent neisserial colonization in humans. Although *N. musculi* does not cause disease, it does encode many pathogenic *Neisseria*-host interaction factors important for colonization that can be considered candidate vaccine antigens, such as components of type IV pili, or outer membrane proteins like LctP, a lactate permease^37,38^. In vitro studies have found that type IV pili of *N. gonorrhoeae* are involved in reprogramming the host transcriptional profile and activation of immune signaling pathways^39^. Our immunophenotyping and transcriptomic results were obtained in the context of an upper respiratory tract long-term persistence model influenced by likely similar antigenic factors. The fact that human pathogenic species like *N. gonorrhoeae* also colonize related tissue like the nasopharynx makes our data on the consequences of oral colonization by *N. musculi* very relatable and significant. Additional pathogenic neisserial antigens could be engineered into *N. musculi* and tested in our model for their effect on interactions with the host and ensuing immune responses. Identification of safe and effective immune profiles and transcriptome changes induced by infection have been used as surrogates for “ideal” vaccine-elicited responses^40^. Similarly, our new model and associated data will be a useful tool for characterizing the in vivo transcriptional and immunological endpoints of *Neisseria*-host mucosal colonization and would likely be useful for development of vaccine candidates against neisserial pathogens. This model system could be used to test the efficacy of *N. musculi* or gonococcal antigen vaccination on mucosal persistence by *N. musculi* or *N. musculi* strains serving as expression vectors for antigens from *Neisseria* species pathogenic to humans. The model could also study how enhancement or suppression of host immune responses influences *N. musculi*-induced innate or adaptive immune responses’ that impact vaccine efficacy against persistent asymptomatic carriage.

## Methods

### Bacterial strains and growth conditions

A naturally occurring RifR smooth morphotype, strain NW742, of *Neisseria musculi*, was used for oral inoculations^41^. *Neisseria musculi* was cultivated on Gonococcal Base (GCB; Difco) agar plates containing rifampin (40 mg/L) and Kellogg’s supplements routinely incubated at 37°C with 5% CO_2_ for 48 hours. 48-hour old colonies were spread onto GCB plates, grown for 18 hours, then suspended in Phosphate Buffered Saline (PBS) and adjusted to an optical density at 600 nm of 1.0 for oral inoculations of mice.

### Mouse inoculations

Sixteen female A/J mice, five weeks old, procured from the Jackson Laboratory (Bar harbor, ME) were acclimatized for one week within the animal facility at Ohio University. Two days prior to *N. musculi* inoculation, oral swabs and fecal pellets were collected to screen for pre-existing *Neisseria* flora. No *Neisseria* spp. were detected after 48 hours of incubation. Mice were evenly divided into two groups of eight mice each, consisting of an inoculated group and a control group. Mice in the inoculated group were manually restrained, and a slow oral inoculation of 50μL bacterial suspension was administered, while the control group were mock inoculated with 50μL of PBS into the oral cavity.

### Confirmation and Sample collection

Three days post-inoculation, oral swabs and fecal pellets were collected and subsequently plated as described earlier to confirm the colonization of *N. musculi* in the inoculated group, with no detection of bacteria in the control group. Following confirmation, on the fifth day, four mice each from both the inoculated and control groups were humanely euthanized using CO_2_ asphyxiation. Blood samples, lungs and spleens were then collected for flowcytometry analysis (described below). Blood collection was conducted through cardiac puncture using a 20-gauge hypodermic needle (BD), resulting in the collection of approximately 2 mL of blood, which was placed into a vacutainer tube. Lungs and spleens were harvested by making dorsal incisions, and the respective organs were carefully extracted using sterile forceps. The collected organs were each placed in a tube with organ transport medium (RPMI with 10% fetal bovine serum) containing penicillin and streptomycin and transported to the Ohio State University on wet ice. The tongues of mice were harvested as previously described^41^ by making incisions made on both sides of the mouth of euthanized mice. The oral cavity was opened, and the tongue was held with sterile forceps from the tip and cut at the base with sterile scissors. Each tongue was cut into small pieces, homogenized with a Mini Beadbeater-16 using 2.3 mm Zirconia beads (BioSpec Products, Bartlesville, OK), diluted in Hanks’ Balanced Salt Solution with 0.01 mM HEPES and 0.3% w/v bovine serum albumin, and plated to enumerate colony forming units (CFUs) per gram of tissue. The hard palate was collected and 1/3^rd^ of each hard palate was homogenized as above and plated for CFU enumeration. The remaining 2/3^rd^ was placed in RNA protect (0.5 mL/palate) and snap-frozen in liquid nitrogen.

### Final Sampling

The remaining eight mice were monitored for 33 days. Weekly fecal and oral swabs were collected to quantify the colonization burden in the inoculated group and confirm the absence of *Neisseria* in the control group. Oral swabs were collected as described previously^41^ and suspended in GCB + 20% glycerol, vortexed for one minute, and serially diluted in GC broth, for plating on GCB agar plates containing rifampin. Fecal pellets were collected in sterile microfuge tubes, weighed, and each pellet was mixed with 1 mL of GCB + 20% glycerol using a sterile plain wooden applicator. The pellet suspensions were vortexed for 1 minute and then plated on GCB agar plates containing rifampin to enumerate CFUs per gram. After 33 days, the remaining mice were euthanized using CO_2_ asphyxiation. Blood samples, lungs, spleens, tongues, and hard palates were collected as described above.

### Ethical Approval

All animal protocols were approved by the Ohio University Institutional Animal Care and Use Committee prior to the initiation of the experiments

### RNA extraction and sequencing

Mouse palates were removed from RNA Protect, rinsed with cold PBS, then subjected to bead beating using 2-3 3mm glass beads on a TIssueLyser II (4 minutes each side, 20Hz) in 350μL of buffer RLT supplemented with 1% betamercaptoethanol and 20ng of cRNA as a carrier. Debris were pelleted and discarded, and total RNAs were extracted from the supernatant with the Qiagen Micro RNeasy kit. RNAs were quantitated using a Bioanalyzer, then ribosomal RNAs depleted using the NEBNext^®^ rRNA Depletion Kit with bacterial and human/mouse/rat probes mixed at a ratio of 30%/70%, respectively. Strand-specific, dual unique indexed libraries were made using the NEBNext^®^ Ultra™ II Directional RNA Library Prep Kit for Illumina^®^ (New England Biolabs, Ipswich, MA). Manufacturer protocol was modified by diluting adapter 1:30 and using 3μL of this dilution. Library size selection was performed with AMPure SPRI-select beads (Beckman Coulter Genomics, Danvers, MA). Glycosylase digestion of adapter and 2^nd^ strand was done in the same reaction as the final amplification. Libraries were QC’ed using the DNA High Sensitivity Assay on the LabChip GX Touch (Perkin Elmer, Waltham, MA). Library concentrations were also assessed by qPCR using the KAPA Library Quantification Kit (Complete, Universal) (Kapa Biosystems, Woburn, MA). Pooled libraries were sequenced on an Illumina NovaSeq 6000 using 150bp PE reads (Illumina, San Diego, CA). Reads were mapped to the *Mus musculus* GRCm39 genome using HISAT^42^.

### RNA-seq data analysis

Gene expression counts were estimated using HTseq^43^. Normalized count data and Variance Stabilized Transformation (VST) counts were generated using the R DESeq2 package^44^. For DEG estimation, inoculated samples were compared to their respective control samples using DESeq2 with an FDR ≤0.05 and an absolute Log_2_ Fold Change ≥1 (“Significant” DEGs, Supplementary Table 1). A heatmap (Supplementary Figure 1) and a volcano plot of immunological biomarkers were generated based on Z-scores of VST counts (Figure 2A). GO analysis was performed using DEGs (with a reduced significance cutoff of p-value ≤ 0.01) with the R package ClusterProfiler v4.0, and Cnet plots were generated using the cnetplot function^45^. All FASTQ files and associated metadata are uploaded to the Gene Expression Omnibus (GEO) repository (see Data availability).

### Deconvolution of transcriptome data

The TIMER2.0 database^12^ was utilized to perform deconvolution of bulk transcriptome data to explore the differential expression of immune cell subsets within tissue colonized by *N. musculi*. Using six algorithms (TIMER, CIBERSORT-ABS, QUANTISEQ, XCELL, EPIC, MMCPCOUNTER) we generated differential estimations of abundance of immune subsets from colonized palate tissue of inoculated mice at day 5 and day 33 of sacrifice and that of control mice. Statistical comparisons using abundance scores of immune cells were performed using the Wilcoxon rank-sum test (p<0.05).

### Tissue processing for flow cytometry staining

Tissue processing and flow cytometry staining were performed as previously described^46^. Briefly, mice were euthanized, and their lungs, spleens, and blood were processed. Tissues were filtered, suspended, and centrifuged. Following lysis and washing steps, cells were resuspended in R-10 media. 100μL of whole blood was stained for flow cytometry using the below protocol. The cells were then fixed with 2% paraformaldehyde and filtered into polystyrene FACS tubes using filter caps.

### Flow staining and data acquisition

Flow cytometry staining and data acquisition were conducted using previously described methods^47^. In brief, splenic and lung tissue samples were divided into two equal portions (1 million cells/mL each). One portion was stimulated with a cocktail of phorbol 12-myristate 13-acetate (PMA) and ionomycin, while the other remained unstimulated. Stimulation included Golgi-Plug and Golgi-Stop, followed by a 20-hour incubation. After fixation and permeabilization, cells were stained and analyzed on a Cytek Aurora flow cytometer. Data were processed using FlowJo v.10.6.2, with “fluorescence minus 1” controls for each marker.

### Graphing and statistics

Graphs were prepared using Graph Pad Prism (version 9.3.1). FlowSOM-based automatic clustering algorithms and Uniform Manifold Approximation and Projection (UMAP) for dimensional reduction visualization were performed using OMIQ. Data was statistically analyzed using their bundled software. Comparisons between two groups were performed using the Wilcoxon rank-sum test. Pearson correlation coefficient analysis was used for correlations between immune signatures and tissue colony counts.

## Data Availability

The datasets generated during and/or analyzed during the current study are available from the corresponding author on reasonable request.

## Acknowledgments

We thank members of the Maryland Genomics Core at the Institute for Genome Sciences (IGS), University of Maryland School of Medicine, for palate sample processing and sequencing. This study was partially supported by Ohio University’s Infectious and Tropical Disease Institute and Molecular and Cellular Biology Graduate program.

## Author Contributions

M.A. and M.G. conducted immunological assessments, analyzed data, and led the overall manuscript writing. T.Z. and S.B. conducted mouse colonization studies and tissue harvests. A.D. performed transcriptome analysis and contributed to manuscript writing. W.M., L.T., and T.S. were involved in tissue processing and flow cytometry experiments. S.W., S.S., and A.W. conducted deconvolution data analysis and contributed to manuscript writing. D.K. supervised data analysis.

H.T. designed the transcriptome analysis and contributed to manuscript writing. N.W. designed the bacterial inoculation study and contributed to manuscript writing. N.L. designed the immunological assessment, integrated microbiological and transcriptome data, and led the overall manuscript writing.

