## Supplemental Figure 1 for "Unveiling the Immune Dynamics of Neisseria Persistent Oral Colonization: A Roadmap for Innovative Vaccine Strategies"

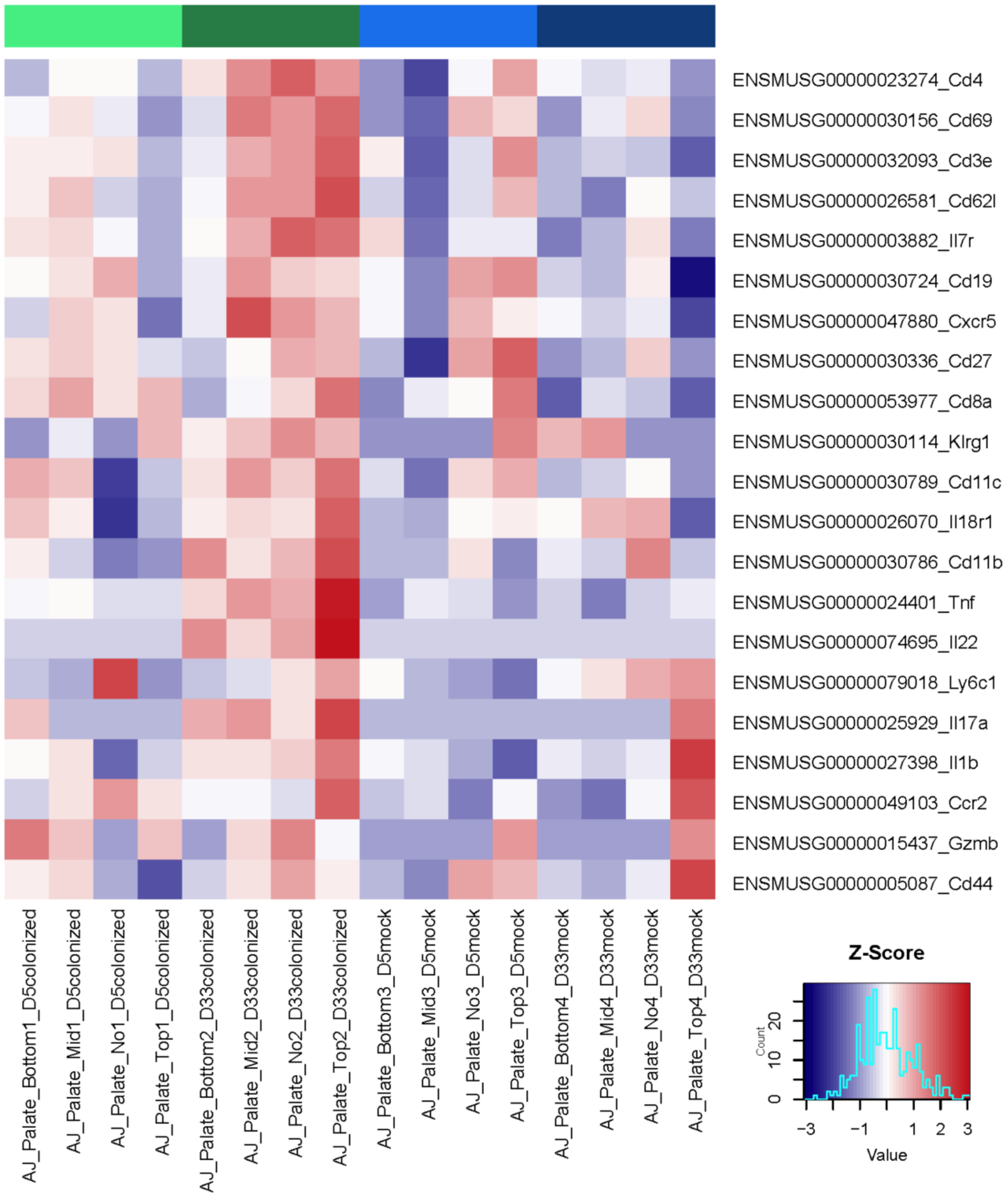


**Supplementary Figure 1.** Heatmap of selected immunological biomarkers was generated based on Z-scores of VST counts. “Colonized” refers to mice inoculated with *N. musculi*. “Mock” refers to, uninoculated control mice.

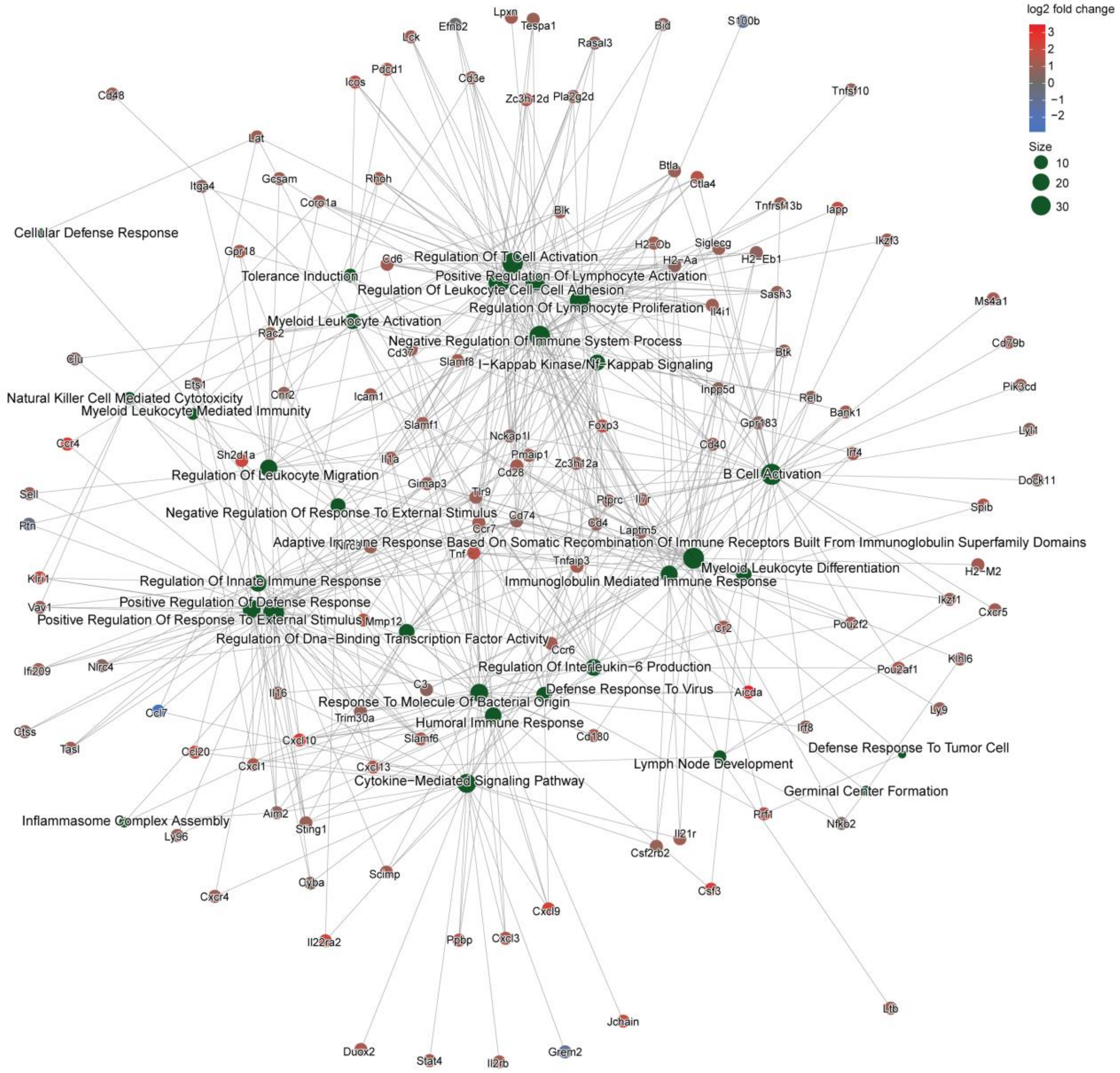


**Supplementary Figure 2.** Expanded Cnet plot of GO biological process enrichment of DEGs of palate tissue of inoculated vs uninoculated mice shows distinct connections between biological processes resulting from colonization with *N. musculi.*

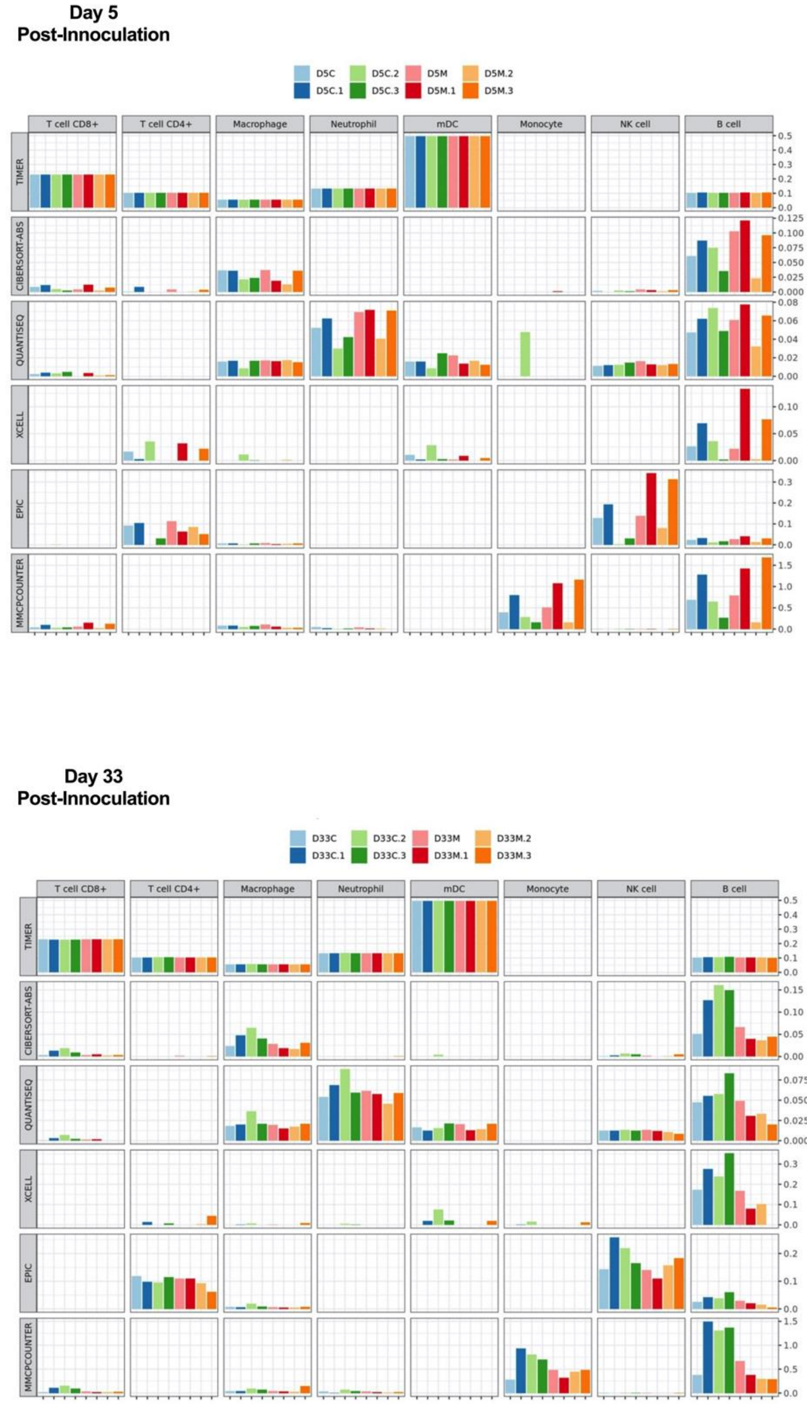


**Supplementary Figure 3.** Deconvolution of palate transcriptome of mice inoculated with *N. musculi* and controls using TIMER2.0 depicted by multi-panel bar plot showing differences of estimations of abundance by six different algorithms among samples.

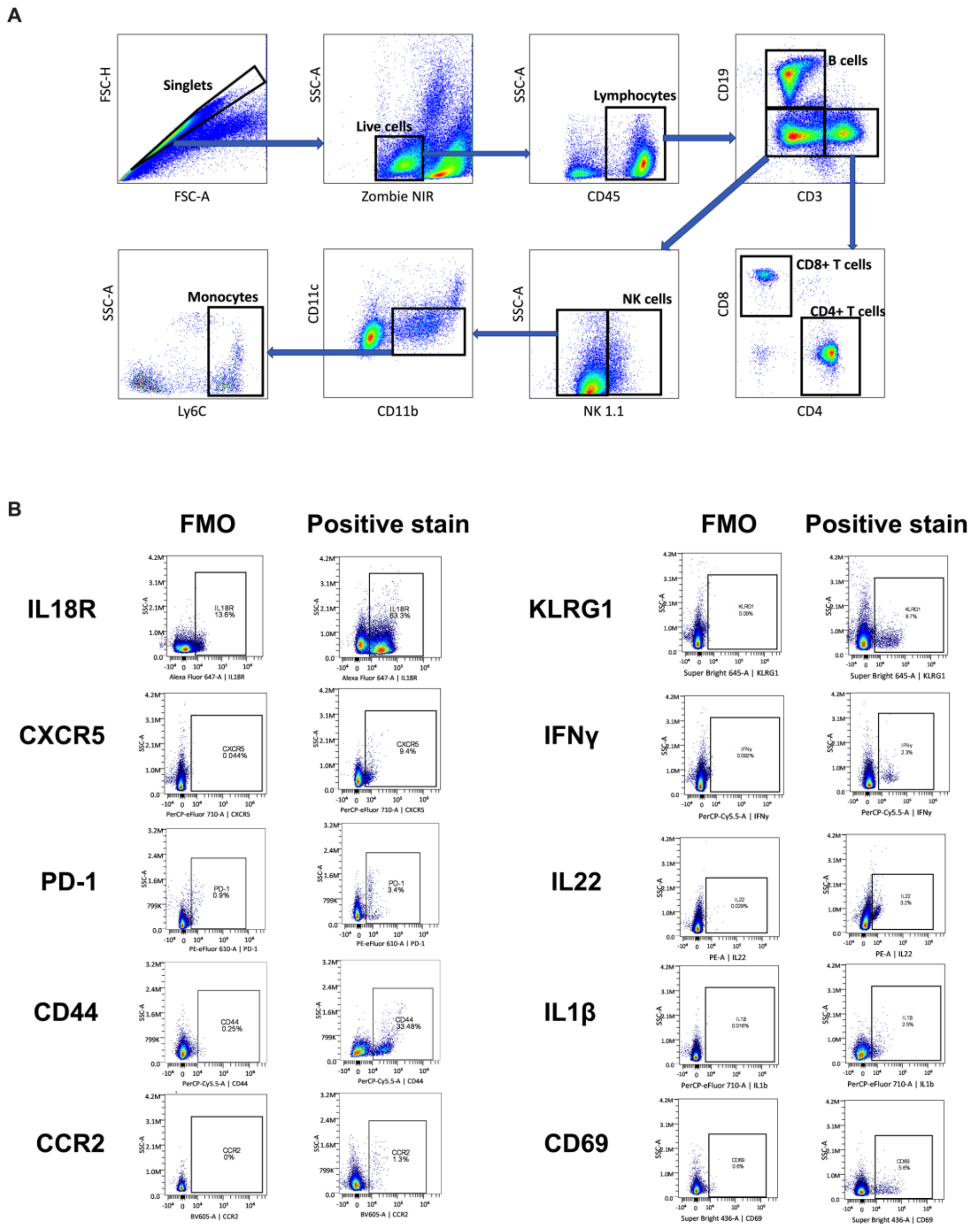


**Supplementary Figure 4. (A)** Representative gating strategy to identify immune cell subsets in analyzed tissue. **(B)** “Fluorescence minus 1” (FMO) controls for each marker prior to spectral unmixing

**Supplementary Table 1. DESeq2 gene lists tested for differential expression between Colonized and Mock infected samples at Day 5 and Day 33, respectively.** Genes were filtered using a False Discovery Rate cutoff of ≤ 0.05 and an absolute $Log_2$ Fold Change cutoff of ≥ 1. Genes that met these cutoffs were classified as "Significant".

**(Attached as excel file)**

**Supplementary Table 2. Complete list of Gene Ontology (GO) terms obtained using differentially expressed genes in palates of mice 33 days after being orally inoculated with *N. musculi* vs uninoculated controls**

**(Attached as excel file)**
